# Control of Cell Division Orientation by Cell-intrinsic and Tissue-scale Forces During *Drosophila* Axis Elongation

**DOI:** 10.64898/2026.06.18.733192

**Authors:** Chloe A. Kuebler, Adam C. Paré

## Abstract

Oriented cell division (OCD) within the plane of a tissue is a conserved mechanism of tissue elongation, maintenance and organization. Here, we investigate how myosin II polarity downstream of leucine-rich-repeat (LRR) receptors influence OCD in the early *Drosophila* embryo. Following convergent extension, cells in the ventral mesectoderm (VME) and cells along compartment boundaries in the lateral neuroectoderm (LNE) divide parallel to the anterior-posterior axis. We analyzed division angles in live mutant embryos lacking myosin polarity at specific cell-cell junctions and show that both cell-intrinsic polarity and non-autonomous outside forces influence division angles differently between the two tissues. We demonstrate that a minimum threshold of myosin polarity, not overall myosin levels, is required to bias divisions as the embryo elongates. In the LNE, we present evidence that strong myosin polarity mediated by Tartan at compartment boundaries biases boundary divisions while simultaneously buffering cells inside compartments from pulling by the posterior midgut. In the VME, our results suggest that signaling by Tartan and Toll family receptors (Toll-2, Toll-6, Toll-8) primes all cells to become responsive to posterior pulling to strongly bias cell divisions. Together, this study shows that cell-intrinsic LRR receptor signaling, primarily mediated by Tartan, interacts with cell-extrinsic pulling forces to shape the distinct patterns of OCD observed in the *Drosophila* germband.

## INTRODUCTION

An important mechanism for properly integrating dividing cells into complex tissues is oriented cell division (OCD), a phenomenon in which daughter cells preferentially segregate along a directional axis. Asymmetric OCD along the apical-basal axis, particularly in the context of the stem-cell niche, has been extensively studied (Quyn et al., 2010; Yadlapalli and Yamashita, 2012). By contrast, symmetric OCD can also occur within the plane of an epithelial tissue, with cells predominantly dividing along a major body axis. During animal development, reproducible patterns of planar OCD are necessary to yield proper cell identity and tissue morphology, and defects in this process have been linked to microcephaly, Huntington’s disease, and kidney malformation (di Pietro et al., 2016; Fish et al., 2006; Noatynska et al., 2012; Saburi et al., 2008; Scepanovic and Fernandez-Gonzalez, 2018). Therefore, there is considerable interest in identifying the molecular cues and biomechanical inputs that control OCD within the plane of a tissue.

The *Drosophila* germband is a tractable model system for studying planar OCDs because specific populations of cells divide in a stereotyped spatial and temporal manner within a short window of developmental time. Embryonic cells on the surface of the germband are separated into two distinct tissues: a narrow band of cells along the ventral midline known as the ventral mesectoderm (VME), and two large populations of cells on either side of the embryo making up the lateral neuroectoderm (LNE)(Foe, 1989). Following gastrulation (stage 6), LNE cells undergo cell intercalation to drive convergent extension of the anterior-posterior (AP) axis (stage 7). Towards the end of convergent extension, all VME cells divide once in a highly biased manner along the AP axis (Camuglia et al., 2022; Wang et al., 2017), and in the LNE, a wave of cell divisions moves from the dorsal to the ventral side of the embryo (stage 8)(Foe, 1989). Previous studies have characterized OCD in these tissues, and shown that both planar polarized MyoII activity and external pulling forces can affect cell division angle (Blanchard et al., 2024; da Silva and Vincent, 2007; Scarpa et al., 2018; Wang et al., 2017). Importantly, the patterned cell-surface molecules that directly control MyoII polarity in distinct regions of the germband are now known (Paré et al., 2019, 2014; Sharrock et al., 2022; Urbano et al., 2018), making it possible to specifically test how cell-intrinsic contraction in specific subsets of cells integrate with cell-extrinsic pulling forces to determine patterns of OCD.

In this study, we set out to determine the relative importance of moderately polarized non-muscle myosin II (MyoII) activity within germband compartments (mediated by Toll family receptors) compared with strongly polarized MyoII activity at compartment boundaries (CBs) (mediated by Tartan). When we ubiquitously increased MyoII activity in an unpolarized manner, division angles exhibited no directional bias, whereas increasing MyoII activity in a polarized manner led to more AP-biased cell divisions across the germband, indicating that OCD requires a threshold level of polarized MyoII activity. Loss of Toll-mediated and Tartan-mediated MyoII polarity led to the randomization of OCDs in the VME, suggesting relatively cell-autonomous effects in this tissue. By contrast, in the LNE, loss of Toll-mediated MyoII polarity had relatively little effect, whereas loss of Tartan-mediated MyoII polarity led to more AP-biased cell divisions. By comparing these effects to a mutant background lacking both cell-intrinsic MyoII polarity and pulling forces from the posterior midgut, we conclude that the role of CBs on OCD in the LNE is twofold. First, high MyoII planar polarity in CB cells leads to cell divisions parallel to the AP axis in a cell autonomous manner. Second, CBs counteract the poster pulling force leading to indiscriminate cell division angles within embryonic compartments. These results demonstrate that autonomous molecular cues, specifically LRR receptors, coordinate cell divisions in conjunction with tissue-scale pulling forces during tissue growth in the early *Drosophila* embryo.

## RESULTS

### Cell divisions are AP-biased in the VME and along CBs in the LNE

To characterize early OCDs in the VME (mitotic domain 14) and LNE (mitotic domain 11), we measured the angle of division (0° parallel to the AP axis) in the middle region of the germband over 30 min (t0 = first cell division in the VME) in wild-type embryos expressing a cell membrane marker (spider–GFP) (Figure 1A,B). The VME begins as two rows of cells located along the ventral midline, and each VME cell divides once towards the end of convergent extension (stage 7/8) (Foe, 1989; Kearney et al., 2004). Cell divisions in the VME were highly biased along the AP axis (median=15.6°) (Figure 1C), resulting in an average VME width of 2.5 cells after divisions were complete (Figure 1B, right). To test whether CBs (Figure 1A) affect OCDs in the VME, we compared division angles between cells adjacent to a CB (CB cells) and non-CB cells. We observed no significant difference in division angle between CB and non-CB cells in the VME during this period (Figure 1C). In the LNE, a wave of cell divisions occurs in a dorsal-to-ventral direction, with the dorsal-most divisions roughly coinciding with the first VME divisions (Figure 1B) (Foe, 1989). LNE cells displayed a slight AP bias (median=39.2°) (Figure 1D). However, when we compared division angles between CB and non-CB cells in the LNE, we found that CB cells were significantly more likely to divide along the AP axis compared with non-CB cells (median=25.3° vs. 42.4°; p=0.016) (Figure 1D). These data demonstrate that different populations of germband cells display distinct patterns of OCD in wild-type embryos, with VME cells and LNE cells at CBs showing an AP-bias, and non-CB cells in the LNE dividing randomly.

**Figure 1.**
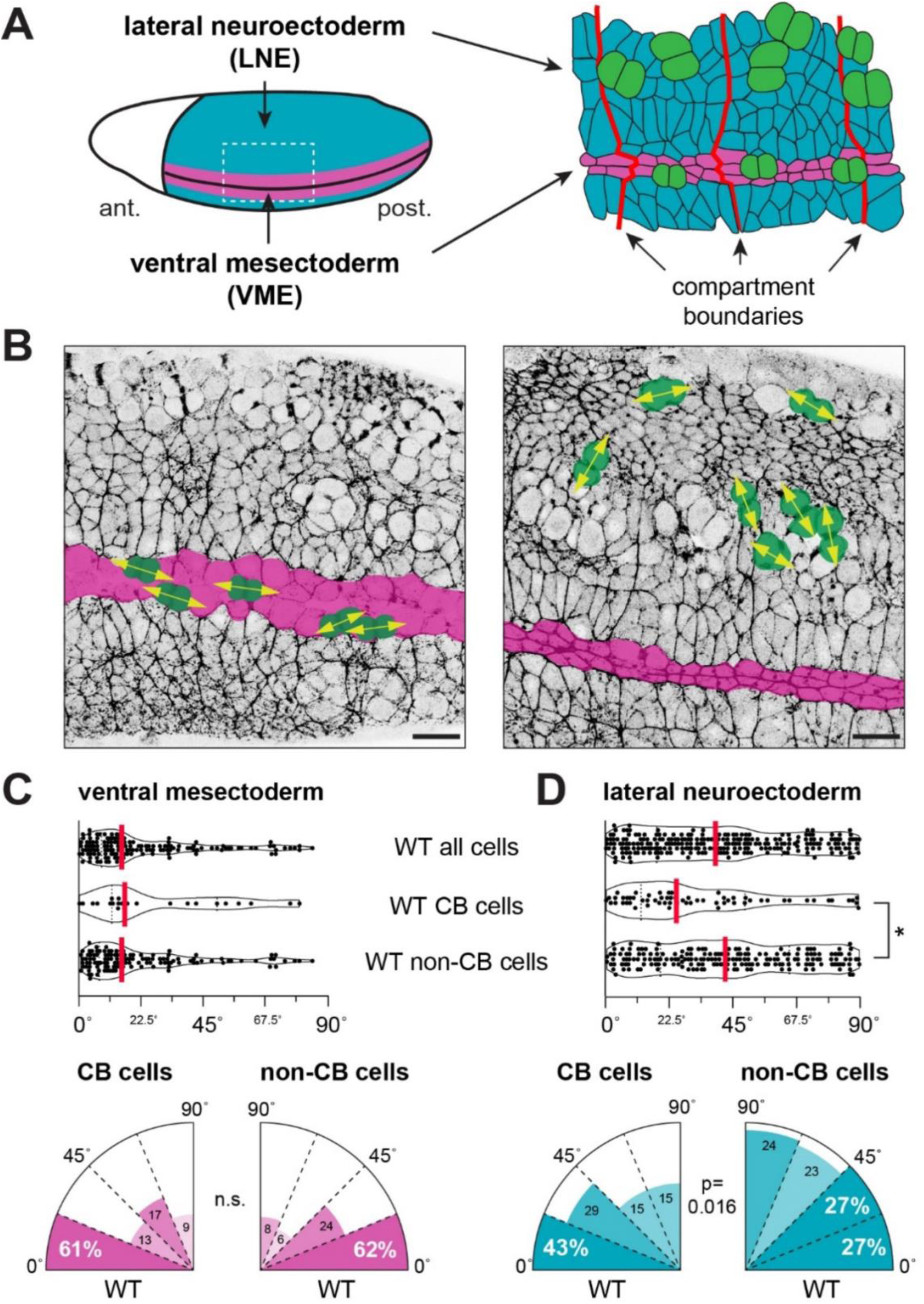
Characterization of oriented cell divisions in the germband epithelium. **A)** The *Drosophila* germband epithelium consists of the lateral neuroectoderm (LNE, blue) and ventral mesectoderm (VME, magenta). Compartment boundaries are shown in red, and dividing cells are shown in green. **B)** Stills from time-lapse movies showing MyoII LC-GFP in control (*sqh–GFP)* embryos; dividing cells are highlighted with yellow arrows in the VME (left) and LNE (right). Scale bars (lower right corner), 20 µm. **C,D)** Violin plots (top) and rose plots (bottom) showing cell division angles in the VME (C) and LNE (D) for all cells in WT (n=148 and 252 cells, respectively) and WT CB (n= 23 and 61 cells, respectively) and non-CB cells (n= 125 and 191 cells, respectively) (K-S test; median, red).

### Increasing MyoII planar polarity induces more AP-biased cell divisions

To test how MyoII activity affects OCDs in the VME and LNE, we used genetic methods to ubiquitously increase MyoII activity in the embryo. Initially, we used the GAL4/UAS system to overexpress the ROK activator ShroomA, which physically tethers ROK to the cell cortex to promote MyoII contractility at adherens junctions (De Matos Simões et al., 2014). If polarized MyoII is responsible for OCDs in the germband, then we hypothesized that overexpressing ShroomA activity might lead to more AP-biased cell divisions, particularly in the LNE. However, when we measured division angles in ShroomA^OE^ embryos, we found that cell divisions had no directional bias compared with overexpression controls in both the LNE (median=39.7° vs. 26.1°; p=0.003) and the VME (median=23.3° vs. 12.5°; p=0.002) (Figure 2A,B). Considering this result contradicted our initial hypothesis, we decided to quantify MyoII levels and planar polarity in ShroomA^OE^ embryos immediately before the onset of cell divisions. In overexpression control embryos, as expected, MyoII LC-mCherry was more highly enriched at vertical (DV-oriented) compared with horizontal (AP-oriented) junctions (median log2(V/H)=0.75) (Figure 2C-E). However, in ShroomA^OE^ embryos, MyoII-mCherry levels were significantly increased at both vertical and horizontal junctions compared with controls (p=0.02 and p<0.0001, respectively) (Figure 2C,D), leading to significantly decreased MyoII planar polarity (median log2(V/H)=0.18) (p<0.0001) (Figure 2E). These results indicate that increasing MyoII levels in an unpolarized manner disrupts OCDs in both the VME and LNE.

**Figure 2.**
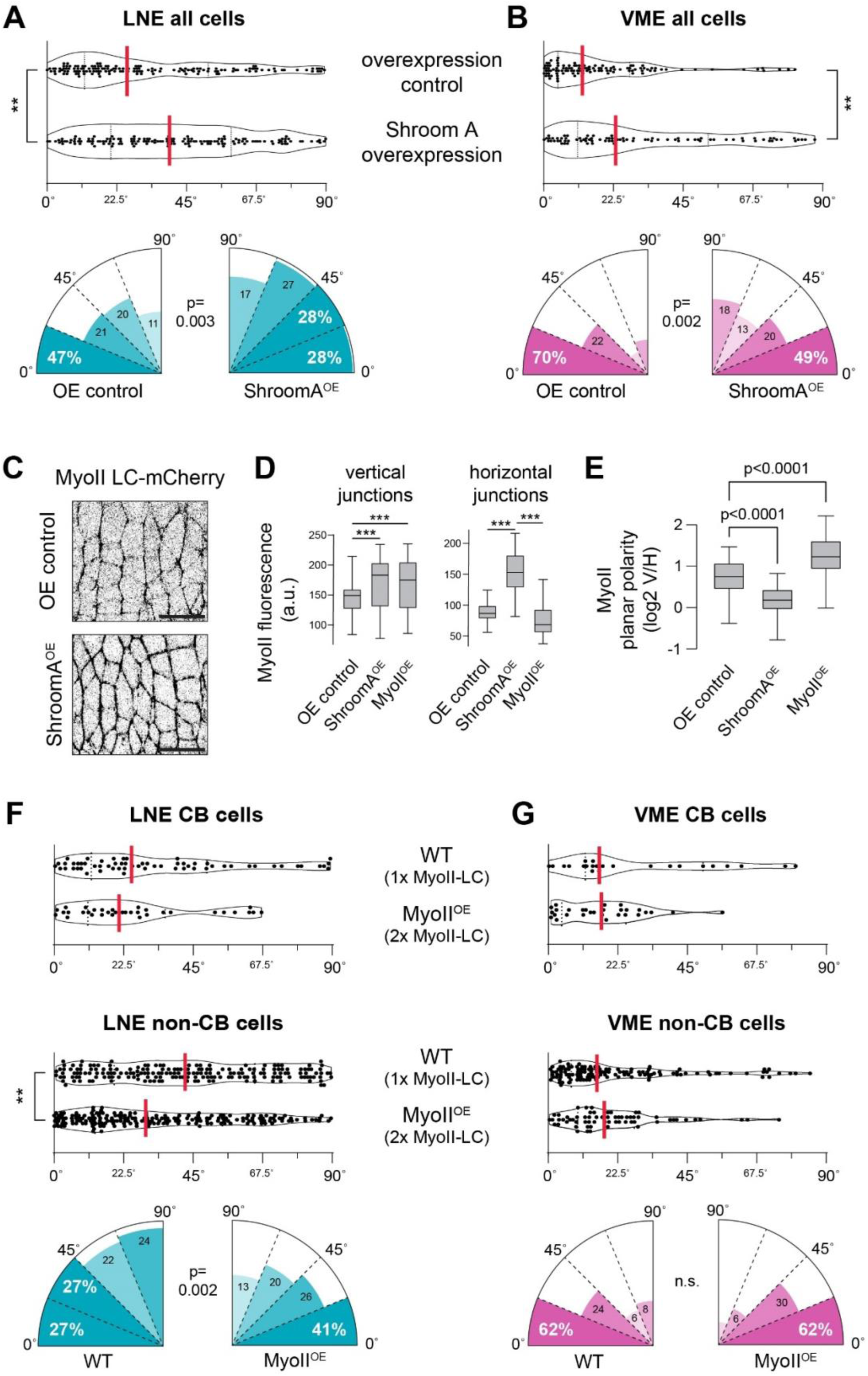
Enhanced MyoII activity influences division angle in the LNE and VME. **A,B)** Violin plots (top) and rose plots (bottom) showing cell division angles in the LNE (A) and VME (B) in controls (n=149 and 112 cells, respectively) and ShroomA^OE^ (n=148 and 84 cells, respectively) (K-S test; median, red). **C)** Stills from time-lapse movies showing MyoII-LC*–mCherry* in control embryos (*mat67–Gal4; 95-1–GFP, sqh–mCherry*), and embryos overexpressing ShroomA (*UAS–ShroomA; mat67–Gal4; 95-1–GFP, sqh–mCherry*). Scale bars (lower right corner), 20 µm. **D)** Quantification of MyoII fluorescence at vertical junctions (left) and horizontal junctions (right) **E)** Quantification of MyoII planar polarity in control, ShroomA^OE^, and MyoII^OE^ embryos. MyoII intensity calculated as a ratio of vertical:horizontal cell interfaces (30 polarity ratios per OE group, 3 embryos each; 40 polarity ratios for controls, 4 embryos) one-way ANOVA. **F,G)** Violin plots (top) showing cell division angles in the LNE (E) and VME (F) in MyoII^OE^ CB (n= 34 and 29 cells, respectively) and non-CB (n=200 and 64 cells, respectively) compared to WT CB (n= 61 and 23 cells, respectively) and non-CB cells (n= 191 and 125 cells, respectively) (K-S test; median, red). And rose plots (bottom) showing non-CB cell division angles.

In the previous experiment, we noticed that cell divisions in the LNE were more AP-biased in the overexpression controls compared with wild-type embryos (median=26.1° vs. 39.2°) (compare Figures 1D and 2A). These “wild-type” embryos had just a maternally provided cell membrane marker, whereas the overexpression control embryos had both a cell membrane marker and a fluorescently tagged version of MyoII light chain (MyoII LC-mCherry). Therefore, we hypothesized that this extra copy of maternally provided MyoII LC might be affecting OCD by increasing MyoII activity in a polarized manner. To test this, we quantified MyoII planar polarity and OCD in a separate stock that carried two extra copies of MyoII LC–GFP, which we refer to as MyoII^OE^. Indeed, we found that MyoII planar polarity was significantly higher in MyoII^OE^ embryos than in the overexpression control (log2(V/H)=1.22 vs. 0.75; p<0.0001) (Figure D,E), and we observed significantly more divisions parallel to the AP axis in MyoII^OE^ LNE non-CB cells compared with wild-type non-CB cells (29.6° vs. 42.4°; p=0.002) (Figure 2E). These effects were specific to LNE non-CB cells, as LNE CB and VME cells were not significantly affected (Figure 2F,G). These results show that increasing planar polarized MyoII leads to more AP-oriented cell divisions specifically within compartments, and they also suggest that OCDs require a minimum threshold of planar polarized MyoII.

### LRR receptor expression patterns change during cell division

MyoII is enriched at vertical junctions in the germband due to the striped expression of four leucine-rich repeat (LRR)-containing cell-surface proteins: Toll-2, Toll-6, Toll-8 and Tartan (Paré et al., 2019). The expression patterns of these four LRR genes prior to convergent extension (early stage 7) have been well characterized in the LNE (Paré et al., 2019, 2014), but how these patterns change over the course of cell division has not been systematically described. We used fluorescent in situ hybridization (FISH) to detect the mRNA expression patterns for these LRR genes directly before the onset of cell divisions (mid stage 7) (Figure 3A-D); *wingless* was used as a spatial reference. We confirmed that the Toll gene expression patterns were similar between the LNE and VME (Figure 3A-C). We note that *Toll-8* showed an additional horizontal band of moderate expression in the ventral LNE and VME cells. Using immunofluorescence, we also verified that Tartan protein is present in alternating tissue compartments in the VME at this stage (Figure 3D). We conclude that germband cells in the LNE and VME display differential expression of all four LRR proteins immediately prior to cell division, consistent with a potential role in mediating OCD.

**Figure 3.**
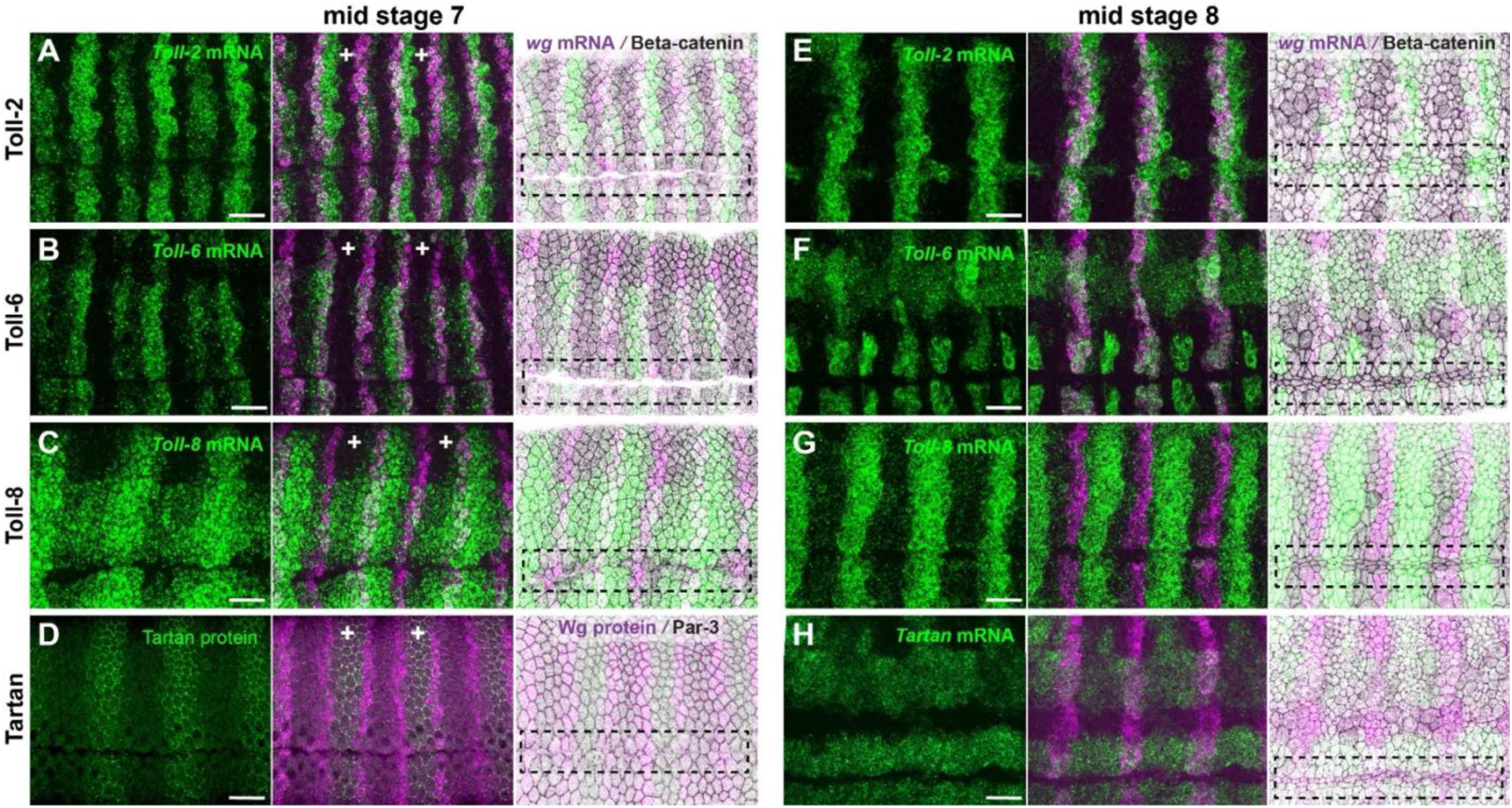
LRR encoding genes that influence MyoII polarity undergo dynamic changes in expression during germband elongation. **A)** mRNA expression patterns of Toll-2, Toll-6, Toll-8, and wingless with Tartan protein expression in stage 7 embryos. Corresponding embryonic compartments indicated by white crosses. In each row, the first panel shows the expression profile of each LRR-encoding gene via *in situ* hybridization (scale bars, 20 µm), the middle panel shows the LRR receptor relative to segment polarity gene *wingless*, and the third panel shows the combination of LRR-encoding gene (green), *wingless* (magenta), and cell membranes (gray). Dashed boxes separate the VME from the LNE. **B)** mRNA expression patterns of Toll-2, Toll-6, and Toll-8, Tartan, and Wingless in stage 8 embryos after VME cells have divided.

We also characterized the expression patterns of the *Toll-2, -6, -8* and *tartan* genes after VME divisions were complete (mid stage) (Figure 3E-H). At this point, compartments have transitioned from ∼4 to ∼8 cells wide due to cell intercalation, and the wave of cell divisions in the LNE has reached the ventrolateral germband. For *Toll-*2, the vertical stripes of expression in the LNE maintained their register and stripe expression levels became more consistent; in the VME, the borders of each *Toll-2* stripe expanded posteriorly by roughly two cells (Figure 3E). For *Toll-6*, the LNE expression pattern became more complex, with a horizontal band of expression appearing in the lateral germband, and vertical stripes transitioning to small patches of expression directly adjacent to the VME; in the VME, *Toll-6* appeared to be completely downregulated (Figure 3F). For *Toll-*8, expression was similar in the LNE and VME, and the pattern was overall less complex, with the horizontal band of expression disappearing and the vertical stripes becoming more distinct (Figure 3G). Finally, Tartan showed the most significant change in expression pattern. Most notably, the vertical stripes of Tartan protein were no longer detectable at the membrane by antibody staining (data not shown), and FISH revealed a complete shift towards horizontal stripes of expression; in the VME, *tartan* appeared to be completely downregulated (Figure 3H). We conclude that the LRR gene expression patterns are quite dynamic during cell division, and it appears that only *Toll-2* and *Toll-8* remain differentially expressed in the VME after divisions are complete.

### Toll receptors influence VME morphology

In the LNE, Toll-2, Toll-6 and Toll-8 are required to establish MyoII planar polarity within tissue compartments and drive cell intercalation during stage 7 (Paré et al., 2019, 2014). To test whether Toll receptors are required for OCD during stage 8, we characterized VME width and cell division angles in *Toll-2,6,8* triple-mutant embryos. We found that final VME width was significantly different between *Toll-2,6,8* and wild-type embryos (3.5 cells vs 2.5 cells; p<0.0001) (Figure 4A). When we analyzed OCD in the VME between *Toll-2,6,8* embryos and wild-type, we observed consistent shifts towards more randomized division angles in the mutants (Figure 4B,C), consistent with a thicker final VME width. This effect was most apparent when considering all cells in the VME (22.6 ° vs. 15.6°), although it only reached borderline significance (p=0.07) (Figure 4B). In the LNE, we also saw a trend towards more randomized cell division angles in the *Toll-2,6,8* embryos compared with wild-type, although these differences did not reach statistical significance (Figure 4D). Taken together, these results show that loss of Toll receptor activity does not significantly affect cell division angles in the LNE, although it does lead to a significantly wider VME morphology, which could be due to subtle defects in OCD.

**Figure 4.**
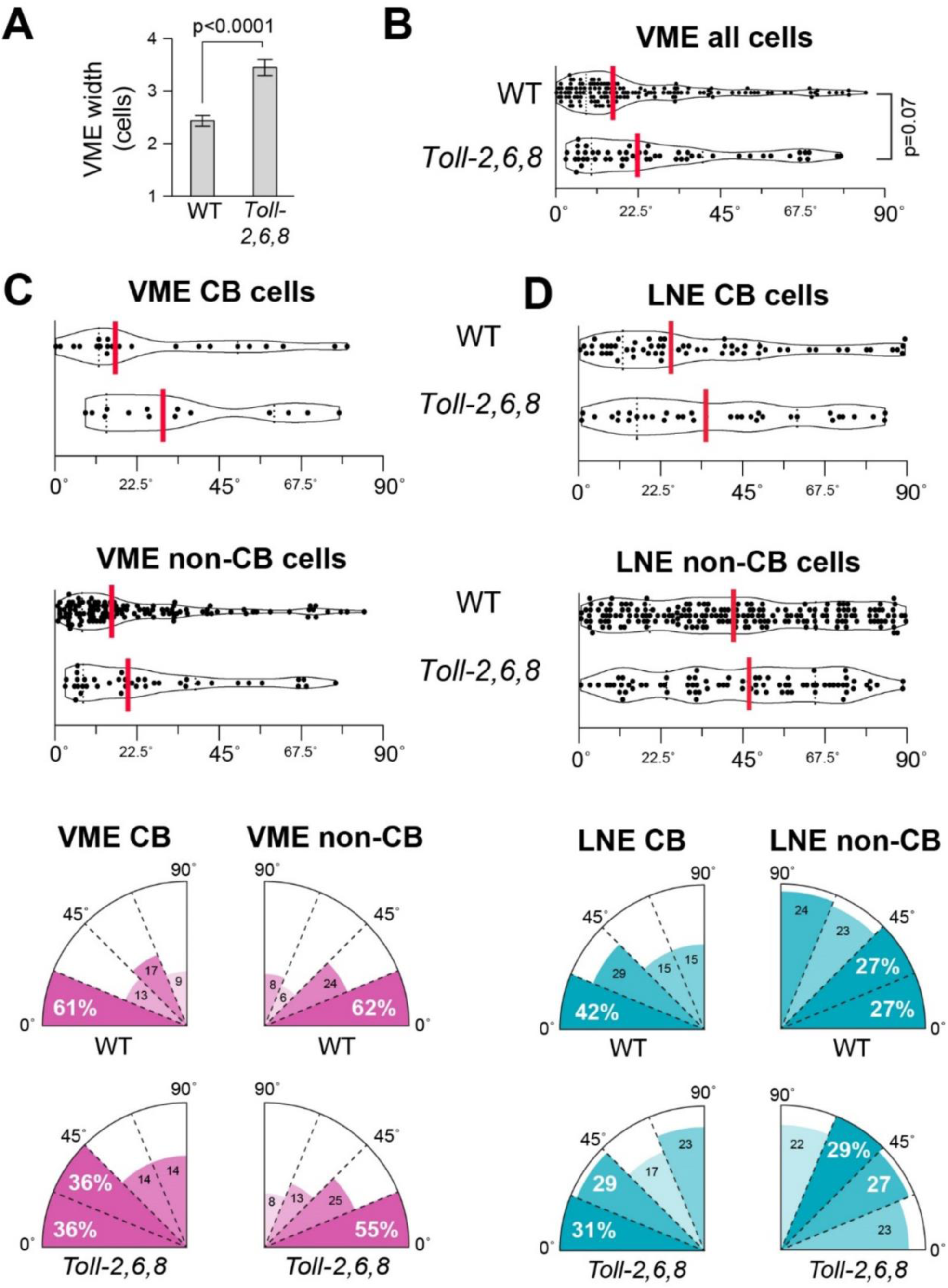
Toll-receptors influence VME morphology. **A)** Histograms of VME width (mean + s.e.m.) after cell divisions in WT (*95-1–GFP*) (n=40 measurements from 4 embryos) and *Toll-2,6,8* embryos (n=30 measurements from 3 embryos) (one-way ANOVA). **B)** Violin plot showing the distribution of all VME division angles in WT (*95-1–GFP,* n=148 cells) and *Toll-2,6,8* (*Toll-2; Toll-6, Toll-8, 95-1–GFP*, n=54 cells) embryos. **C,D)** Violin plots (top) and rose plots (bottom) showing division angles in the VME (C) and LNE (D) in Toll-2,6,8 CB (n= 14 and 35 cells, respectively) and non-CB (n=40 and 79 cells, respectively) compared to WT CB (n= 23 and 61 cells, respectively) and non-CB cells (n= 125 and 191 cells, respectively) (K-S test; median, red).

### Loss of Tartan affects OCD in distinct ways between the VME and LNE

Both Tartan and Wingless have been shown to be necessary for different aspects of CB formation in the germband (Paré et al., 2019; Sharrock et al., 2022; Urbano et al., 2018). However, it is not clear whether CBs affect cell behaviors in the VME, or if their roles are restricted to the LNE. To test this, we characterized final VME width and OCD in *tartan* and *wingless* single-mutant embryos. We observed a highly significant randomization of division angles in the VME between *tartan* mutants and wild type (median=31.1° vs. 15.6°; p<0.0001) (Figure 5A,C). Consistent with this, we also saw a significantly greater final VME width in *tartan* mutants (3.3 cells vs. 2.5 cells; p<0.0001) (Figure 5D). By contrast, *wingless* mutant embryos were not significantly different than wild type with respect to VME division angle (Figure 5A) or final VME width (Figure 5D). These data demonstrate that Tartan activity is necessary to orient cell divisions in the VME, whereas Wingless appears dispensable at this stage.

**Figure 5.**
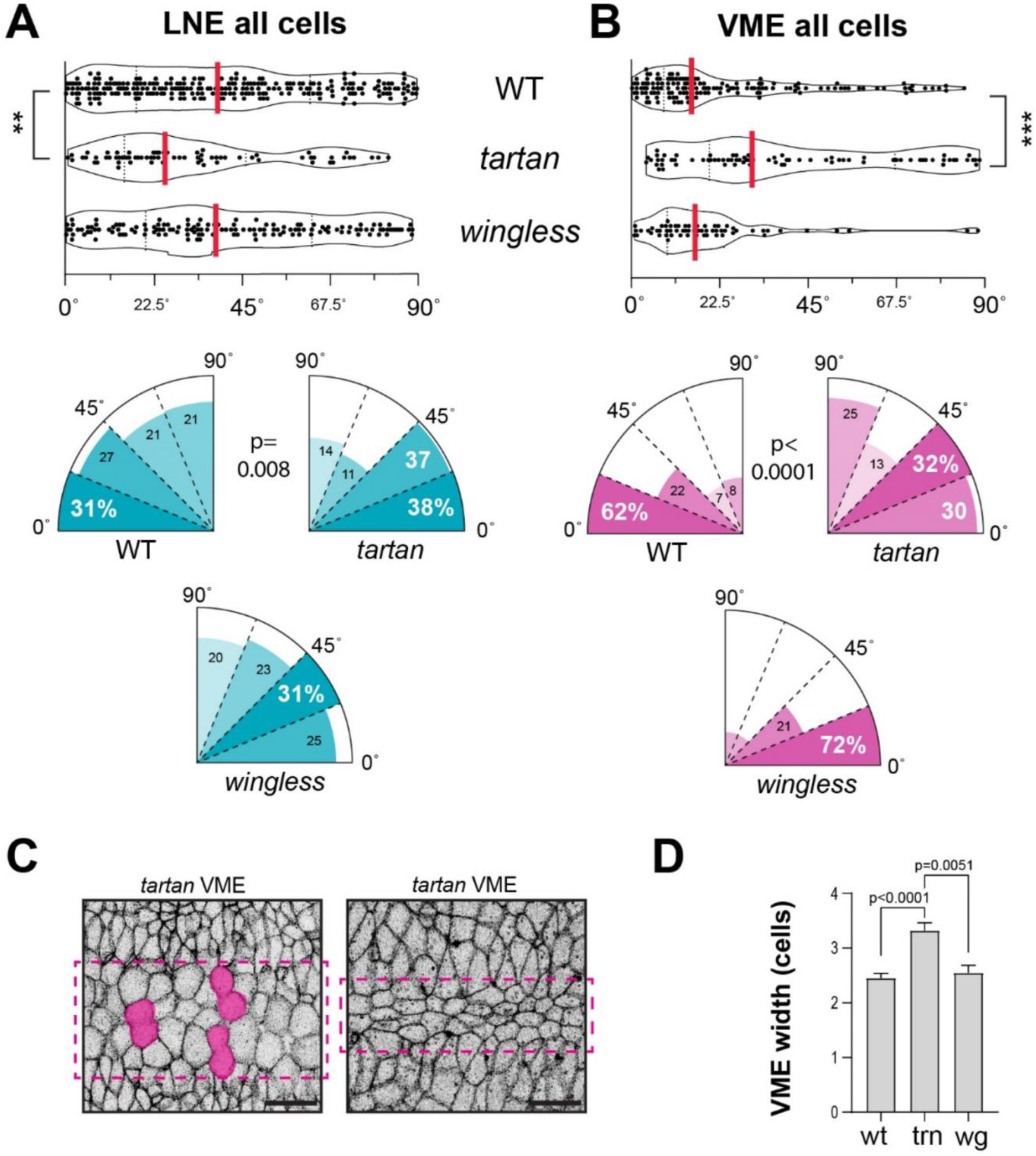
Tartan establishes compartment boundaries and influences OCDs across the germband. **A,B)** Violin plots (top) and rose plots (bottom) showing division angles in the LNE (A) and VME (B) in *tartan* (n= 73 and 77 cells, respectively) and *wingless* (n=143 and 61 cells, respectively) embryos, compared to WT (n=252 and 148 cells respectively) (K-S test; median, red). **C)** Still images from *tartan* (*tartan, 95-1–GFP*) mutant embryos during VME divisions (left)and after VME divisions (right); highlighting VME cells (magenta) dividing perpendicular to the AP axis and final VME width. Dashed boxes indicate VME cells (scale bars, 20 µm). **D)** Histograms of VME width (mean + s.e.m.) after cell divisions in WT (*95-1–GFP*) (n=40 measurements from 4 embryos) and *tartan* embryos (n=30 measurements from 3 embryos) (one-way ANOVA).

Considering that CB cells divide parallel to the AP axis in the LNE, we hypothesized that we would see changes in OCD in this tissue if we disrupted CB formation. Therefore, we also tested the effects of *tartan* and *wingless* mutations on OCD in the LNE. Because there are no CBs in *tartan* embryos at this stage, we were only able to compare all LNE cells between the genetic backgrounds. We did indeed observe a highly significant difference in division angles across all cells in the LNE between *tartan* and wild type (median=25.5° vs. 39.2°; p<0.0001). However, this shift was in the opposite direction to what we expected, meaning loss of *tartan* led to more, rather than fewer, AP-biased cell divisions in the LNE. By contrast, loss of *wingless* had no effect on OCD in the LNE. These results indicate that Tartan-mediated CBs are necessary for the proper distribution of cell division angles in the LNE, and they also demonstrate that loss of *tartan* affects OCD in opposite ways in the VME and LNE.

### Compartment boundary polarity and extrinsic pulling forces contribute to OCDs

Considering the loss of *tartan* randomizes cell divisions in the VME but leads to more AP-biased divisions in the LNE, this indicates that a direct relationship between MyoII planar polarity and AP-oriented divisions within individual cells cannot fully explain the pattern of OCDs in the germband. Another source of directional information that germband cells might be using to orient cell divisions is the posterior pulling force exerted on the germband by posterior-midgut invagination. To test the importance of this external force, we characterized OCD in embryos injected with H1152, a highly specific inhibitor of Rho kinase (ROK), the direct activator of MyoII (Kerridge et al., 2016; Sasaki et al., 2002). Embryos were injected near the cephalic furrow to disrupt MyoII polarity in the anterior germband without disrupting posterior pulling forces from midgut invagination. Live imaging showed a robust loss of cortical MyoII–GFP in ROK inhibitor embryos compared with water-injected controls (Figure 6A). In the VME, division angles were essentially randomized in ROK inhibitor embryos compared with controls (median=52.7° vs. 25.2°; p=0.001) (Figure 6B). By contrast, in the LNE, divisions were skewed along the AP axis in ROK inhibitor embryos compared with controls (median=21.7° vs. 28.8°; n.s.) (Figure 6C), reminiscent of what was seen in *tartan* mutant embryos. We note that very few cell divisions were observed in the LNE in these experiments, perhaps because cytokinesis is more sensitive to ROK inhibition in the LNE, which made it difficult to collect sufficient data points to demonstrate a significant difference. Nevertheless, these data are consistent with the idea that OCDs in the LNE and VME are differentially affected by the loss of MyoII activity.

**Figure 6.**
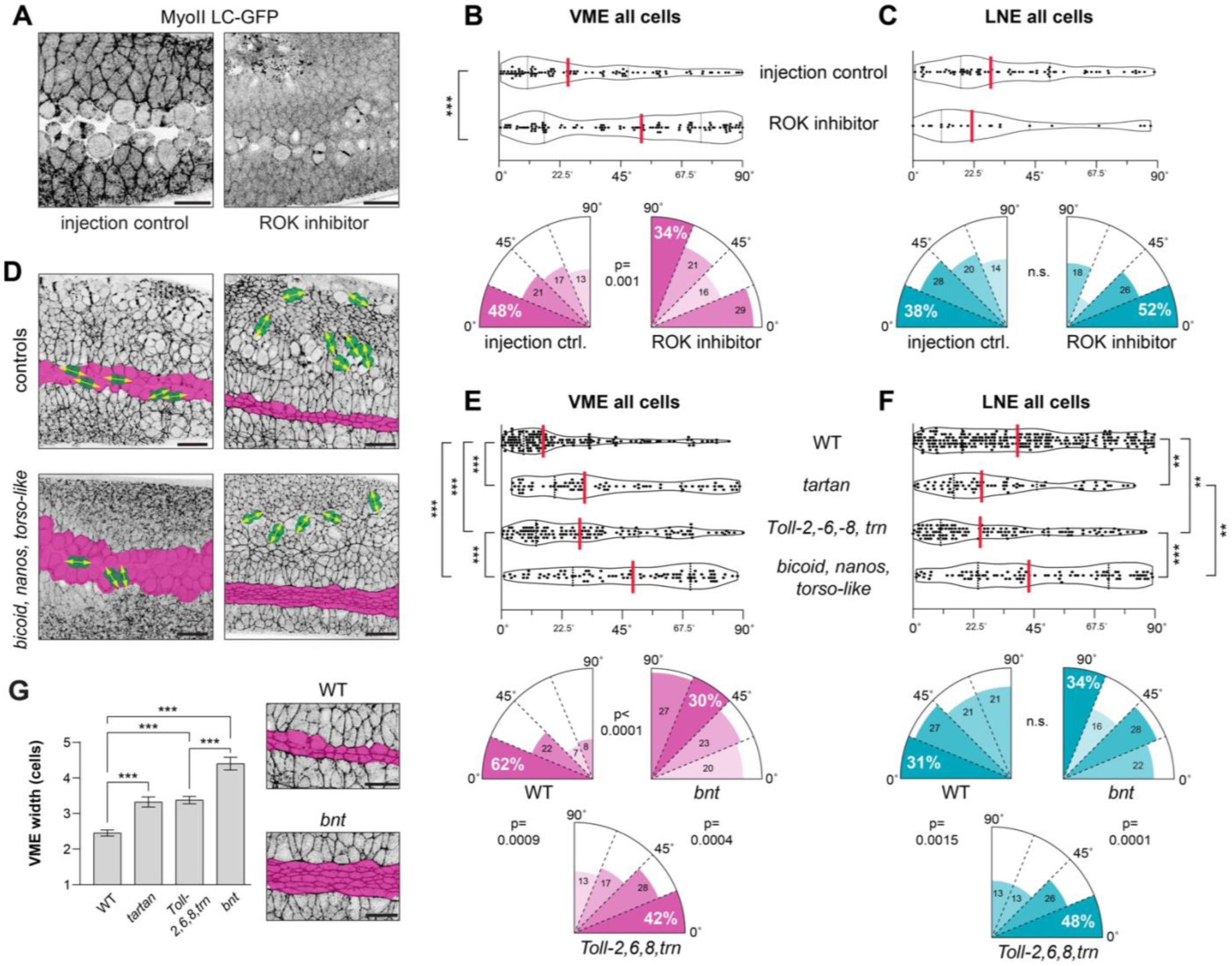
Eliminating CB MyoII polarity and terminal patterning skews division orientation. **A)** Stills from time-lapse movies showing MyoII-LC; images from water-injected control embryo (*gap43–mCherry, sqh–GFP*, 3 embryos) (top), and embryo injected with 5 µM ROK inhibitor, H1152 (*gap43–mCherry, sqh–GFP*, 3 embryos) (bottom). **B,C)** Violin plots (top) and rose plots (bottom) showing division angles in the VME (B) and LNE (C) for all cells in control (n=98 and 76 cells, respectively) and ROK inhibitor-injected embryos (n=114 and 23 cells, respectively). **D)** Stills from time-lapse movies showing MyoII LC-GFP in control (*sqh–GFP)* and *bnt* (*bicoid*, *nanos*, *torso-like* maternal mutants with *sqh–GFP*) embryos; dividing cells are highlighted with yellow arrows in the VME (left) and LNE (right). Scale bars (lower right corner), 20 µm. **E,F)** Violin plots (top) and rose plots (bottom) showing division angles for all VME (E) and LNE (F) cells in WT (n=148 and 252 cells, respectively), *tartan* (n= 77 and 73 cells, respectively), *Toll-2,6,8,trn* (n=125 and 118 cells, respectively), and *bnt* (n=88 and 89 cells, respectively) embryos (K-S test; median, red). **G)** Histograms of VME width (mean + s.e.m.) after cell divisions in WT (*95-1–GFP*) (n=40 measurements from 4 embryos), *trn* (n=30 measurements from 3 embryos), *Toll-2,6,8,trn (Toll-2; Toll-6, Toll-8, tartan, 95-1–GFP)* embryos (n=30 measurements from 3 embryos), and *bnt* (n=30 measurements from 3 embryos) embryos (one-way ANOVA). Representative stills from WT and *bnt* embryos showing final VME morphology (right) scale bars, 20 µm.

To genetically test the influence of this posterior pulling force, we compared OCD in *Toll-2, Toll-6, Toll-8, tartan* quadruple-mutant (*Toll-2,6,8,trn*) embryos and *bicoid, nanos, torso-like* (*bnt*) mutant embryos. *Toll-2,6,8,trn* mutants lack nearly all planar polarized MyoII in the trunk of the embryo but still undergo posterior midgut invagination (Paré et al., 2019). *bnt* embryos lack both planar polarized myosin and posterior midgut invagination, due to the absence of AP patterning and terminal signaling, respectively. We hypothesized that if this posterior pulling force does not affect cell division angle, then the patterns of OCD should look similar in these two backgrounds.

In the VME, division angles were quite different between *bnt* and *Toll-2,6,8,trn* embryos (median=48.9° vs. 29.1°; p<0.0001), with the distribution of division angles being more uniform in *bnt* embryos, and divisions retaining a partial AP-bias in *Toll-2,6,8,trn* embryos (Figure 6D,E). We note that the distributions of angles in the *Toll-2,6,8,trn* and tartan single mutants were very similar to one another (n.s.), and both significantly less skewed than wild-type (Figure 6E). *bnt* embryos also had a significantly wider final VME width compared with *Toll-2,6,8,trn* (4.4 vs. 3.4 cells; p<0.0001) (Figure 6G). These results suggest that cell divisions in the VME cells are influenced by Tartan in a cell autonomous manner and by posterior midgut pulling in a non-cell autonomous manner.

In the LNE, division angles were also very different between *bnt* and *Toll-2,6,8,trn* embryos (median=43.3° vs. 24.9°; p=0.002), with divisions again displaying no directional bias in *bnt* embryos (Figure 6D,F). By contrast, LNE divisions in *Toll-2,6,8,trn* embryos adopted a significant AP-bias that closely matched what we saw in *tartan* single mutants, and which was clearly different from the largely random distribution seen in the wild-type LNE. These results indicate that the AP-oriented cell divisions in the LNE observed in *Toll-2,6,8,trn* embryos do not require LRR-mediated MyoII planar polarity and likely occur in response to pulling forces from posterior midgut invagination.

## DISCUSSION

In this study, we set out to better understand how OCDs are controlled during active tissue remodeling by analyzing cell division in two tissues that display distinct patterns of OCD in the *Drosophila* germband. Planar polarized MyoII is critical for many types of cell rearrangements during development, and it has also been demonstrated that planar polarized MyoII can reorient the mitotic spindle independent of cell geometry to directly influence division angle (Blanchard et al., 2024; Lye et al., 2024; Mao et al., 2011; Nakajima et al., 2013). We analyzed embryos early in axis extension (late stage 7/early stage 8), as the germband is becoming partitioned into distinct embryonic compartments through the formation of CBs, continuous lines of vertical cell-cell junctions that are highly enriched for MyoII (Monier et al., 2011; Paré et al., 2019; Tetley et al., 2016). Within compartments, vertical junctions are also enriched for MyoII compared with horizontal junctions, although not to the same degree as at CBs. We found that cells located along CBs displayed a significant AP division bias, whereas non-CB cells within compartments displayed no bias, similar to later stages in this tissue (Scarpa et al., 2018). We show that recruiting excess MyoII to the cortex in an unpolarized manner by overexpressing the ROK activator ShroomA randomizes division angles in the germband. This confirms that planar polarized MyoII activity, and not just high MyoII levels, are required for OCD. We also analyzed OCD in a transgenic background carrying twice the normal levels of MyoII light chain; we observed that LNE cells within compartments, which normally divide randomly, became significantly more likely to divide along the AP axis. By contrast, OCD was not affected by extra copies of MyoII light chain in the VME or in LNE CB cells, two cell populations that already have very high levels of MyoII planar polarity. These results are consistent with a model in which a threshold level of MyoII enrichment at vertical junctions is required to reorient the mitotic spindle to drive horizontal cell divisions.

In the *Drosophila* germband, MyoII becomes planar polarized at the beginning of stage 7 to drive cell intercalation and convergent extension. MyoII is enriched at vertical junctions in response to the striped expression of four leucine-rich repeat (LRR) cell-surface proteins, Toll-2, Toll-6, Toll-8 and Tartan. Together, the Toll receptors generate moderate MyoII planar polarity inside of compartments (Paré et al., 2014), whereas Tartan generates high MyoII polarity specifically at CBs (Paré et al., 2019; Sharrock et al., 2022). We confirmed that these four LRR genes are expressed in non-uniform striped patterns at the end of convergent extension, immediately before the onset of cell divisions (late stage 7), and are thus well positioned to serve as local spatial cues for OCD. Consistent with the idea that a minimum threshold of MyoII polarity is necessary to alter division angle, removing the Toll receptors that mediate relatively weak MyoII polarity within compartments had no significant effect on OCD in the LNE. We did observe distinct effects on OCD in the LNE between *Toll-2,6,8* and *tartan* embryos.

Based on the idea that high Tartan-based MyoII activity at CBs can bias division angles, we hypothesized that we would see a complete randomization of divisions in *tartan* embryos. However, we saw the opposite effect, with divisions across the entire germband becoming more AP-biased in *tartan* single mutants. We also found that the distribution of division angles in quadruple-mutant embryos lacking both Toll and Tartan expression matched the strong AP-bias observed in *tartan* single-mutants. We conclude that Tartan-mediated effects on MyoII polarity at CBs determine the pattern of OCDs in the LNE.

If removing MyoII polarity specifically from CBs––which only represent a small proportion of total cell-cell junctions––can induce cells across the LNE to divide preferentially along the AP axis, this suggests that forces from outside germband can influence OCD (Collinet et al., 2015; Lye et al., 2015). We hypothesize that the role of CBs in the LNE is twofold. First, high MyoII polarity at CBs acts in a cell-autonomous manner to bias division angles along the AP axis. Second, CBs act to dissipate forces from outside the tissue, so that non-CB cells divide randomly, potentially promoting germband bending and overall architecture. One obvious candidate for an outside force that could affect OCD in the LNE is the posterior-directed pulling force from midgut invagination. Indeed, when we analyzed *bnt* embryos, which lack patterned Toll and Tartan expression as well as posterior midgut invagination (Nüsslein-Volhard et al., 1987), we found that division angles in the LNE were completely random. Taken together, these results indicate that CB tension at regular intervals along the AP axis of the embryo buffers non-CB cells from posterior pulling forces and that LNE cells have no division bias in the absence of CB tension and posterior pulling forces.

In the VME, where the vast majority of cells divide parallel to the AP axis, the effects of Toll receptors and Tartan appeared more additive and cell autonomous. In wild-type embryos, the VME was ∼2.5 cells wide after divisions, consistent with a strong AP bias. Whereas in *bnt* embryos the VME ended up being ∼4.5 cells wide, consistent with essentially random divisions. Removal of either the Toll receptors or Tartan both led to an increase in VME width to ∼3.5 cells, suggesting these two signaling systems contribute equally to OCD in the VME. However, when we directly measured division angles in *Toll-2,6,8* mutants, we did not see as strong an effect as in *tartan* single-mutant embryos. Therefore, it is possible that Toll receptors do not strongly affect OCD in the VME, but do affect final VME width, perhaps through other mechanisms such as cell-cell adhesion (Iijima et al., 2020). With respect to the importance of the posterior pulling force on OCD in the VME, laser ablation experiments have shown that cell divisions in the VME become random when the tissue is mechanically separated from posterior-pulling forces (Wang et al., 2017). Therefore, we hypothesize that in wild-type embryos, Tartan (and perhaps Toll receptor) signaling mechanically primes VME cells to become fully responsive to the posterior pulling force. In the absence of posterior pulling, Tartan- and Toll-mediated effects are not by themselves sufficient to induce OCD, so you see random divisions. In the absence of Tartan-mediated polarity, the pulling force is sufficient to induce intermediate OCD in the VME.

The precise molecular mechanisms connecting Tartan-mediated MyoII planar polarity to the mitotic spindle are unclear. Previous studies have shown that the spindle orientation protein, Pins, is planar polarized at vertical interfaces in the VME, and that disrupting Pins polarity randomizes OCDs in this tissue (Camuglia et al., 2022). One possibility is that Pins is polarized in response to patterned Tartan and Toll receptor expression, either directly or indirectly through MyoII activity. Considering our ROK inhibitor injections completely randomized division angles in the VME, which presumably would not affect Tartan localization or expression pattern, this would be consistent with a model in which Tartan polarizes MyoII, which in turn polarizes Pins in the VME. However, Pins in not strongly planar polarized in the LNE (Scarpa et al., 2018), suggesting that the mechanism of action downstream of Tartan is different between these two tissues. Therefore, at CBs in the LNE, it is possible that planar polarized MyoII can influence mitotic spindle orientation directly, as has been proposed (Blanchard et al., 2024), or in some Pins-independent fashion, perhaps involving the interaction partner of Tartan, Ten-m.

## ACKNOWLEDGEMENTS

We would like to thank J. Zallen for providing the Tartan, Par-3 and β-catenin antibodies. We would like to acknowledge generous funding from the NIH NIGMS to A. Paré that directly supported this research (R01GM147372 and R15GM143729), as well as funding and support from the Arkansas Integrative Metabolic Research Center (AIMRC) (P20GM139768), an NIH Center of Biomedical Research Excellence (COBRE). Funding from the Arkansas Biosciences Institute (ABI) helped purchase the Zeiss LSM900 confocal microscope. Stocks obtained from the Bloomington Drosophila Stock Center (NIH P40OD018537) were used in this study. We would also like to acknowledge FlyBase (http://flybase.org).

## METHODS

### *Drosophila* husbandry, stocks, and crosses

Adult flies used to generate embryos were reared on a standard food mixture containing molasses, cornmeal, yeast, malt, agar, and preservatives (Tegosept and propionic acid) and maintained at 25°C in ventilated cages with apple juice/agar plates and yeast paste. Stock lines were maintained between 18–25°C. Embryonic stages used in this study were as follows (at 25°C): stage 6 ≈ 2.5–3.0 h AEL, stage 7 ≈ 3.0–3.25 h AEL, and stage 8 ≈ 3.25–4.0 h AEL. The sex of the embryos was not considered relevant and was not determined in this study.

The following transgenes were used: 1) *95-1–GFP,* transgene to tag apical cell membranes (*Gilgamesh–GFP, aka “Spider–GFP”,* on chromosome III) (Martin et al., 2009). 2) *sqh–GFP,* transgene to tag MyoII (GFP fused to spaghetti squash, the MyoII light chain, driven by the *sqh* promoter on chromosome III (Royou et al., 2002). 3) *sqh–mCherry,* transgene to tag MyoII (mCherry fused to spaghetti squash driven by the *sqh* promoter on chromosome III (Martin et al., 2009). 4) *gap43–mCherry*, transgene to tag cell membranes (mCherry fused to rat Gap43 gene containing a myristoylation sequence) (Martin et al., 2010). 5) *eve–YFP,* transgene to genotype chromosome III in embryos (a bacterial artificial chromosome containing *even-skipped–SFYFP* in its genomic context, reintroduced onto chromosome III) (Paré et al., 2019). 6) *CTG* balancer, to genotype chromosome II in embryos (*CyO* balancer carrying a *twist–GFP* transgene). 7) *mat67-Gal4*, for UAS overexpression in the maternal germline (*αtubulin–Gal4[67]* on chromosome II, gift of Jennifer Zallen). 8) *UAS-ShroomA* (on X chromosome, BDSC #59019).

The following null-mutant chromosomes were used: 1) “*bnt”* chromosome III (*bicoid*, *nanos*, and *torso-like*) (Nüsslein-Volhard et al., 1987). 2) *Toll-2* chromosome II (*Toll-2[attP],* endogenous *Toll-2* ORF replaced with attP-3XP3-DsRed-attP) (Paré et al., 2019). 3) *wingless* chromosome II (*wg[I-8]*, BDSC #5351). 4) *tartan, 95-1–GFP* chromosome III (*trn[Δ3C]*, endogenous *tartan* ORF deleted) (Paré et al., 2019). 5) *Toll-6, Toll-8, 95-1–GFP* chromosome III (*Toll-6[attP],* endogenous *Toll-6* ORF replaced with attP-3XP3-DsRed-attP; *Toll-8[Δ6B]*, endogenous *Toll-8* ORF deleted) (Paré et al., 2019). 6) *Toll-6*, *Toll-8*, *tartan*, *95-1–GFP* chromosome III (same as previous chromosome plus *tartan[Δ3A]*, endogenous *tartan* ORF deleted) (Paré et al., 2019).

### Crosses

**Figure 1**: To generate wild-type (WT) embryos, homozygous *95-1–GFP* virgin females were crossed to *yellow, white* (BDSC #1495) males. F1 heterozygous siblings were allowed to mate, and F2 embryos were imaged. **Figure 2**: To generate embryos overexpressing ShroomA, *mat67-Gal4; 95-1–GFP, sqh–mcherry/TM3* virgin females were crossed to *UAS–ShroomA/Y* males. F1 *UAS–ShroomA/+; mat67–Gal4/+; 95-1–GFP, sqh–mCherry/+* virgin females and F1 *+/Y; mat67–Gal4/+; 95-1–GFP, sqh–mCherry/+* males were selected and crossed, and F2 embryos were imaged. Overexpression control embryos were generated using the same strategy as above, except *UAS–ShroomA/+; mat67–Gal4/+; 95-1–GFP, sqh–mCherry/+* virgin females were crossed to *yellow, white* males. MyoII^OE^ embryos were collected from a *sqh–GFP* (III) homozygous stock. **Figure 4**: To generate *Toll-2,6,8* zygotic triple-mutant embryos, *Toll-2/CyO; eve–YFP* virgin females were crossed to *Sp/CTG; Toll-6, Toll-8, 95-1–GFP/TM6B* males. F1 *Toll-2/CTG; Toll-6, Toll-8, 95-1–GFP/eve–YFP* virgin females and males were selected and crossed, and F2 embryos were imaged. *Toll-2, Toll-6, Toll-8* mutant embryos were identified by a lack of both mesodermal *twist–GFP* and striped *eve–YFP* expression. **Figure 5**: To generate *tartan* zygotic mutant embryos, *Sp/CTG; tartan, 95-1–GFP/TM3,Sb* virgin females were crossed to *Sp/CyO; eve–YFP* males. F1 *Sp/CTG; tartan, 95-1–GFP/eve–YFP* virgin females and males were selected and crossed, and F2 embryos were imaged. *tartan* zygotic mutant embryos were identified by a lack of striped eve-YFP expression. To generate *wingless* zygotic mutant embryos, embryos were collected from a *wg/CTG; 95-1* stock, and homozygous mutant embryos were identified by a lack of *twist–GFP* expression. **Figure 6**: ROK inhibitor and injection control embryos were collected from a *gap43–mCherry;sqh–GFP* (III) homozygous stock. To generate *Toll-2,6,8,tartan* zygotic quadruple-mutant embryos, the same strategy was employed as above to create the Toll triple mutants, except the *Toll-6, Toll-8, tartan, 95-1–GFP* chromosome was used. To generate *bnt* maternal triple-mutant embryos, a *sqh–GFP; bnt/TM3, Sb* stock was used. Homozygous *sqh–GFP; bnt* virgin females and males were selected from the stock and crossed, and F1 embryos were imaged. All embryos from *bnt* homozygous mutant mothers displayed the maternal *bnt* phenotype; zygotic genotypes were not determined.

### Whole mount mRNA detection

To prepare embryos for hybridization chain reaction (HCR) fluorescent in situ hybridization (FISH) plus immunofluorescence, embryos were fixed using a 1:1 solution of 18% formaldehyde in 0.5 µM EGTA in 1x PBS: heptane for 20 min. The vitelline membrane was chemically removed using 100% methanol with vigorous shaking for 30 s. Embryos were washed with 100% xylenes for 15 min and then rehydrated in PBST (PBS with 0.1% Tween-20) for 20 min. Embryos were permeabilized using acetone for 10 min at -20°C, and post-fixed with 5% formaldehyde in PBST for 2 min, followed by multiple PBST rehydration washes. HCR v3.0 split initiator probes were used as described by Choi et al. (2018). Probe sets were designed by Molecular Instruments to target exons present in all gene isoforms for target mRNAs. Embryos were pre-hybridized in probe hybridization buffer for 30 minutes at 37°C before hybridization with 0.8 pmol of each probe overnight at 37°C. Post-hybridization embryos were washed 4 times with probe wash buffer at 37°C, and twice with 5X SSCT before being pre-amplified in amplification buffer for 20 min at room temperature. Embryos were incubated with fluorescent hairpins overnight in amplification buffer at room temperature and then washed 5 times with 5x SSCT (saline-sodium citrate with 0.1% Tween-20) before immunofluorescence staining (below). Hybridization, probe wash, and amplification buffers were as previously described (Choi et al., 2018).

### Immunofluorescence

HCR-stained embryos were blocked in PBS with 10% bovine serum albumin for 30 min at room temperature before incubation with primary antibodies overnight at 4°C. Embryos were washed 4 times with 5x SSCT, blocked again for 30 min at room temperature, and incubated with secondary antibodies for 2 h at room temperature. Finally, embryos were washed 5 times with 5x SSCT before mounting or storage at 4°C.

The following primary antibodies were used: rabbit anti-Tartan (Chang et al., 1993) (1:400), mouse anti-Wingless (DSHB 4D4) (1:250); rabbit anti-β-catenin (Armadillo; gift of Dr. Jennifer Zallen’s lab) (1:150); and guinea pig anti-Par-3 (Bazooka; gift of Dr. Jennifer Zallen’s lab) (1:150). The following secondary antibodies were used at 1:500 dilution: goat anti-mouse Alexa674 (Thermo Fisher #A-21236), goat anti-rabbit Alexa546 (Thermo Fisher #A-11035), and goat anti-guinea pig Alexa568 (Thermo Fisher #A-11075).

### Fixed and live embryo confocal imaging

Fixed and stained embryos were mounted between two coverslips in ProLong™ Gold Antifade Mountant (Invitrogen P36934). Fluorescence signals were collected using an inverted Zeiss LSM900 laser-scanning confocal microscope with a 40x oil-immersion 1.3 N.A. objective. Non-saturated images were collected at optimal confocal resolution and Z-stacks were acquired with 0.5-µm step intervals. Images were processed using Fiji/ImageJ (Schneider et al., 2012).

To collect embryos for live imaging, adults were maintained in ventilated cages with apple-juice agar plates and yeast paste, and allowed to lay for 1–3 h at 25°C. Embryos were dechorionated in 1:1 bleach:water for 3 min, rinsed well, and placed in halocarbon oil 700 (HO700) (Millipore Sigma #H8773) using a fine paintbrush. Gastrula (stage 6) embryos were identified using a dissection microscope and mounted in HO700 between an oxygen-permeable membrane (YSI Life Sciences #66155) and a #1.5 glass coverslip in a custom mounting slide. To track cell divisions, signals from fluorescent transgenes were captured from the ventrolateral apical surface of the embryo. To visualize fluorescent transgenes, laser power and gain were adjusted to collect non-saturated 8-bit images, and stacks were collected every 30 s at 0.7-1x zoom (760-900 x 700-800 pixels) using a 1 AU pinhole and 0.5-µm step intervals between slices. MyoII transgene channels were acquired with 2x averaging.

### Embryo injections

Embryos were collected for 1 h at 25° C, dechorionated in 50% bleach for 3 minutes, immobilized with glue on coverslips, desiccated in a Drierite-containing chamber for ∼10–15 minutes, and then covered in a 1:1 mix of halocarbon oil 700:27. The Rho-kinase pharmacological inhibitor, H1152 dihydrochloride (MedChemExpress, HY-15720A) was injected at a concentration of 5 µM (which we determined have moderate effects) in the perivitelline space around the start of stage 7 (after ventral furrow formation and posterior midgut invagination) and immediately imaged via confocal microscopy, as described above. Water injections were used as negative controls.

### Cell Division Tracking

Using Fiji/ImageJ, apical slices from each timepoint were projected (maximum intensity) to produce the final movies for analysis. Different time-lapse movies were registered based on the first dividing ventral mesectoderm cell in the field of view. Cell division angles were measured manually in projected timelapse movies using Fiji/ImageJ. Angles were averaged across two consecutive frames for actively dividing cells, which were identified by a dumbbell morphology indicative of cytokinesis. Division angles were measured as the angle between a line perpendicular to the cleavage furrow and the ventral furrow (0°).

### Statistical analyses

For division angles, we compared pooled measurement distributions using the Kolmogrov-Smirnov (K-S) test. Within-sample variance was evaluated via One-Way ANOVA and Turkeys multiple comparison tests for each genotype, and no statistical differences between samples were detected. VME width, MyoII fluorescence, and planar polarity log_2_ ratios were evaluated via One-Way ANOVA. Error bars represent s.e.m. unless otherwise specified.

## Notes

### Competing Interest Statement

The authors have declared no competing interest.

